# Mobius Assembly: a versatile framework for Golden Gate assembly

**DOI:** 10.1101/140095

**Authors:** Andreas I. Andreou, Naomi Nakayama

## Abstract

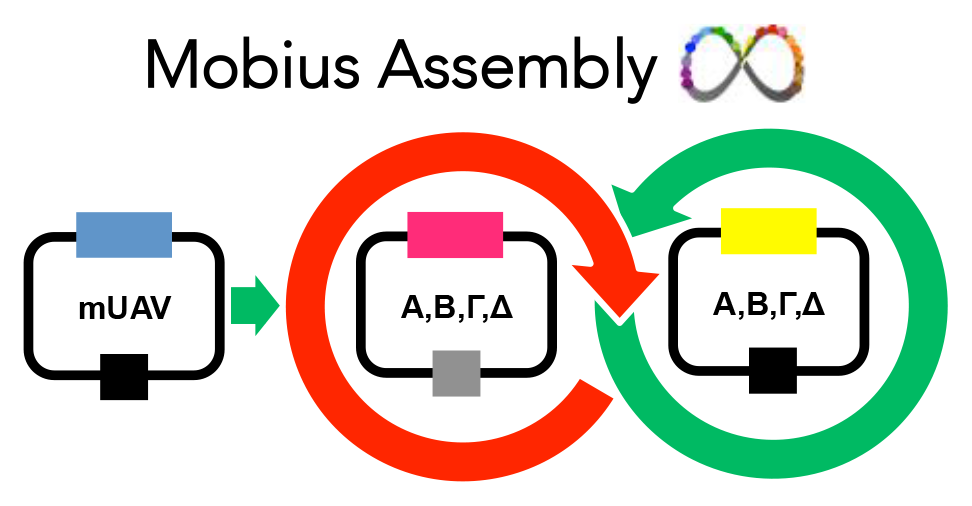

Golden Gate Assembly is a powerful synthetic biology tool, which utilizes Type IIS enzymes for unidirectional assembly of multiple DNA fragments. The simplicity of its DNA assembly and the exchangeability of standard parts greatly facilitate the generation of combinatorial assembly libraries. Currently there are two popular Golden Gate Assembly frameworks that allow multigene augmentation (MoClo and Golden Braid); they render either high cloning capacity or vector toolkit simplicity. We have developed a new Golden Gate Assembly framework called Mobius Assembly, which combines vector toolkit simplicity with high cloning capacity. Mobius Assembly is based on a two-level approach and embraces the standard overhangs defined by MoClo and Golden Braid to confer exchangeability, but with reduced domestication requirement. Furthermore, we have implemented drop-out cassettes encoding chromogenic proteins for visible cloning screening. As proofs of concept, we have functionally assembled up to 16 transcriptional units of various pigmentation genes in both operon and multigene arrangements.

---

Synthetic biology is an interdisciplinary field that adopts engineering principles to facilitate modification of living organisms to provide them with novel functionalities^1^. The engineering principles, such as standardization, modularity, and simplicity, reduce the unpredictability of the living systems, enabling the implementation of the engineering ‘design-construct-test’ cycle in biology. Modular DNA parts (e.g. promoters, coding sequences and terminators) are assembled to build molecular devices (e.g. functional transcriptional units), which can be combined further to assemble genetic modules (e.g. biosynthetic pathways). In addition, standardization sets rules on how these modular parts are designed and assembled^2^. The use of standard parts and assembly methods facilitates exchangeability among users, allowing the reusability of available constructs as well as automatization of construction. Additionally, simple design would aid efficiency and versatility of molecular engineering.

DNA assembly is a pivotal technology in synthetic biology as it materializes the construction of new transcriptional units (TUs) and genetic modules from individual DNA parts^3^. DNA synthesis technologies is rapidly improving to provide affordable *ex vivo* synthesis of large DNA sequences; however, DNA assembly is the predominant strategy for the assembly of DNA fragments larger than 1 kb^4^. Currently, there are several DNA assembly methods used across the synthetic biology community (for an extensive review, see^5^). Those methods mainly fall in three categories: long-overlap assembly (e.g. Gibson Assembly^6^), site-specific recombination that exploits phage integrases (e.g. Gateway cloning^7^), and restriction endonuclease-based strategies. The endonuclease-based assembly methods are the most commonly used category that allows standardization when following specific rules.

One of the first attempts to standardize a restriction enzyme-based DNA assembly method was BioBricks^8^. The reusability and simplicity of BioBricks make them popular, and there have been some efforts to alleviate their drawbacks, such as the in-frame stop codon in the fusion scar and frequent need for ‘domestication’ (i.e. elimination of internal restriction sites^9–11^). However, the pairwise nature of BioBrick assembly renders the construction of multipart systems time-consuming. Cloning is labor-intensive, since the digestion and ligation take part in separate reactions.

Almost a decade ago, a new generation of assembly technology called Golden Gate was introduced^12,13^. It is based on Type IIS restriction endonucleases, which cleave double-stranded DNA outside their recognition sites. They leave a short singlestranded overhang, whose sequence can be defined by users. Golden Gate employs SsaI restriction sites, which are eliminated during subcloning, allowing the simultaneous digestion and ligation in a one-pot reaction. Additionally, the use of distinct 4bp overlaps allows directional, scarless cloning of multiple parts. Nonetheless, the original Golden Gate Assembly framework lacked reusability, since the composite parts (e.g. TUs) could not be assembled further in multigene constructs. It also lacked standardization, since no rule was defined for the 4bp overhangs. Major breakthroughs came when MoClo and Golden Braid variants of Golden Gate Assembly were developed to enable the hierarchical construction of multi-TUs and the full reusability of composite parts^14–17^.

Golden Braid 2.0 framework uses a simple pairwise approach where multipartite expansion is achieved by switching between two levels, α and Ω^15^. The core vector toolkit of Golden Braid is comprised of only five plasmids. To date, the Golden Braid toolkit is mainly targeted at plant systems, although it can be made compatible with other chassis with a few modifications. On the other hand, MoClo framework uses a complex, yet high capacity vector toolkit to achieve parallel assembly^16,17^. The standard parts feed Level 1, which is comprised of seven plasmids. Assembly of up to six TUs takes Level 1 to Level 2 or to Level M/P, each of which involves seven vectors and a suite of End-linkers. The MoClo toolkit was initially released for general eukaryotic expression^16^, which was then adapted for plants^18^, mammalian cells^19^ and yeast^20,21^, and it has just recently been extended for use in E. coli^22–24^ with modifications in the vectors and/or assembly standards.

Speed, simplicity, and capacity are important characteristics for a DNA-assembly method; however, the two most popular Golden Gate variants compromise at least one of them. Golden Braid sacrifices capacity over simplicity, while MoClo emphasizes the capacity of the multi-gene constructs, thus increasing in complexity. A simple assembly method is easier for users to assimilate and to troubleshoot cloning or vector toolkit problems. On the other hand, high capacity assembly methods are preferred, as they are time- and cost-effective. In addition, since both methods implement Type IIS restriction enzymes that are frequent cutters, they are burdened with heavy requirements for domestication, which is labor-intensive.

To address the tradeoffs and limitations of these current methods, we developed Mobius Assembly, a new framework for hierarchical Golden Gate Assembly. Mobius Assembly embodies both simplicity and cloning capacity and thus allows exponential and theoretically unlimited augmentation of TUs. The two-level design, comprised of four Acceptor Vectors in each level and seven auxiliary plasmids, enables a quadruple assembly with a compact vector toolkit. Mobius Assembly also adopts the 4bp standard overhangs defined by MoClo and Golden Braid to promote the sharing of standard parts. Another new feature, the introduction of the rare cutter *Aar*l reduces domestication needs. Furthermore, the vectors are demarcated with specific visible markers for cloning selection. As a proof of concept, we have used Mobius Assembly to successfully reconstruct multi-gene biosynthetic clusters to produce protoviolaceinic acid and carotenoids. Additionally, to validate the capacity of the cloning system, we built a 16TU construct.

## Features of the Mobius Assembly framework

The Mobius Assembly framework commences at Level 0, which represents the standard part library. It uses the Mobius Universal Acceptor Vector (mUAV), to convert amplified PCR fragments into standard, exchangeable parts (Figure 1A). mUAV has a backbone of pSB1C3 and thus confers chloramphenicol resistance. We introduced the chromoprotein amilCP^25^ as a visible cloning selection marker, which imbues a purple colour to the colonies (Figure 2B) (see below for the choice of the visible reporter genes). This negative selection marker is flanked by AarI recognition sites. AarI cuts through the *Bsa*l sequence, generating fusion sites (CTCT and TGAG) where a PCR fragment will be cloned in (Figure 1A). The insert should be amplified with a pair of primers each of which bear an AarI restriction site, a fusion site for the mUAV, and a 4bp standard overhang, from 5’ to 3’. AarI digestion releases the *amilCP* gene, which is replaced by a standard part, resulting in a Level 0 vector. It should be noted that users can use any backbone in all levels of Mobius Assembly if the backbone we provide does not meet specific experimental requirements. To provide this flexibility we flanked the Golden Gate cassettes with *EcoR*l and *Pst*l restriction sites, and the backbone can be swapped with a simple digestion/ligation step.

**Figure 1.**
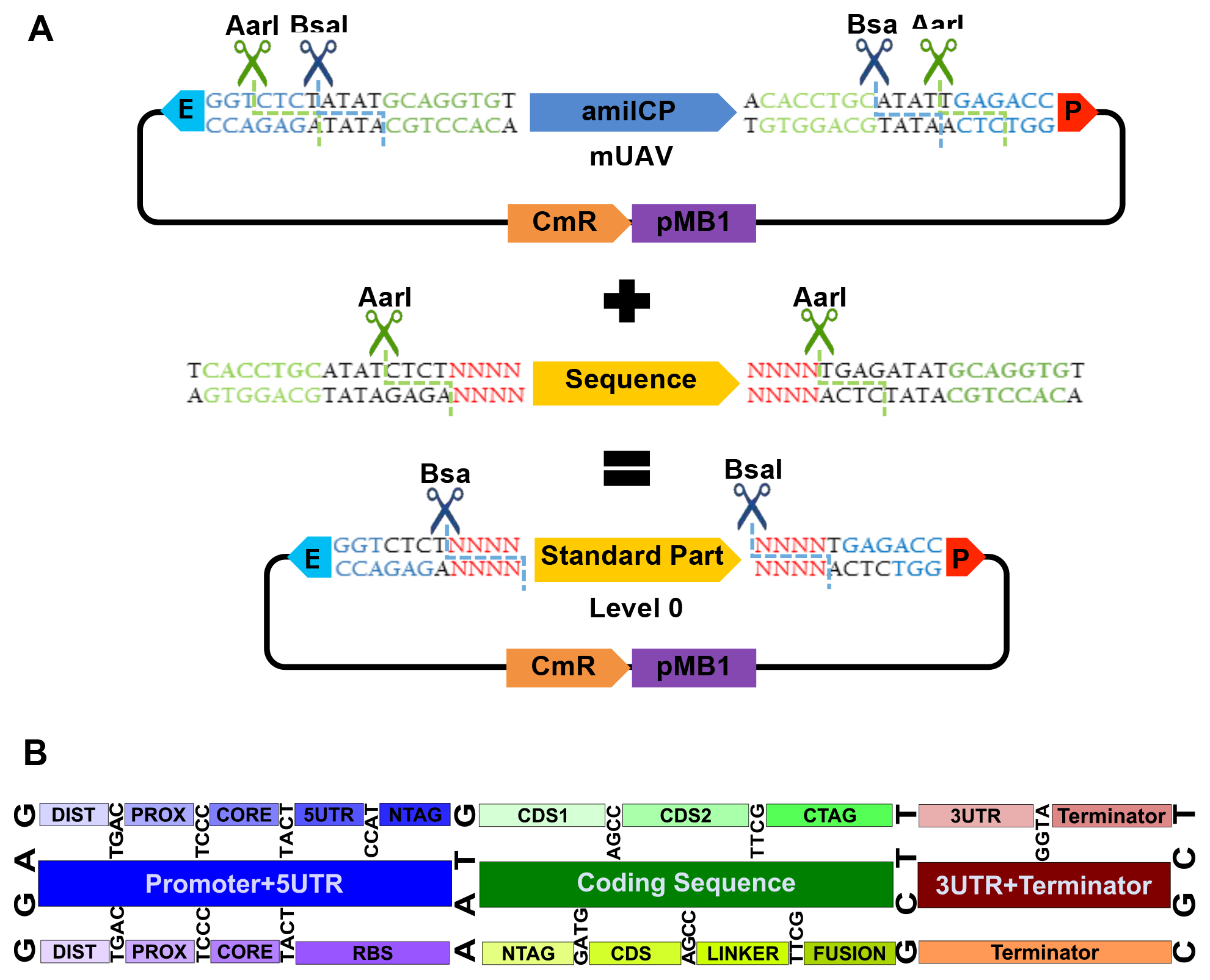
Mobius Assembly standard part generation. (A) Mobius Universal Acceptor Vector (mUAV) is the vector which converts and hosts DNA fragments as standard parts. mUAV is flanked by the Type IIS restriction enzymes *Bsa*l and *Aar*l and carries *amiICP* gene as visible cloning selection marker. The inserts are amplified with primers containing *Aar*l recognition sites, the fusion sites with the mUAV, and the standard overhangs, and they replace *amiICP* cassette in a Golden Gate reaction. The standard parts are released by *Bsa*l digestion. E: *Eco*RI; P: *Pst*l. (B) Mobius Assembly embraces the 4bp standard part overhangs defined by MoClo, Golden Braid, and Phytobricks, to facilitate part sharing. The middle row illustrates the standard overhangs for major functional parts (promoter, coding sequence, and terminator); the top row shows the recommended overhangs for eukaryotic sub-functional parts, while the bottom row indicates ones for the prokaryotic counterparts.

**Figure 2.**
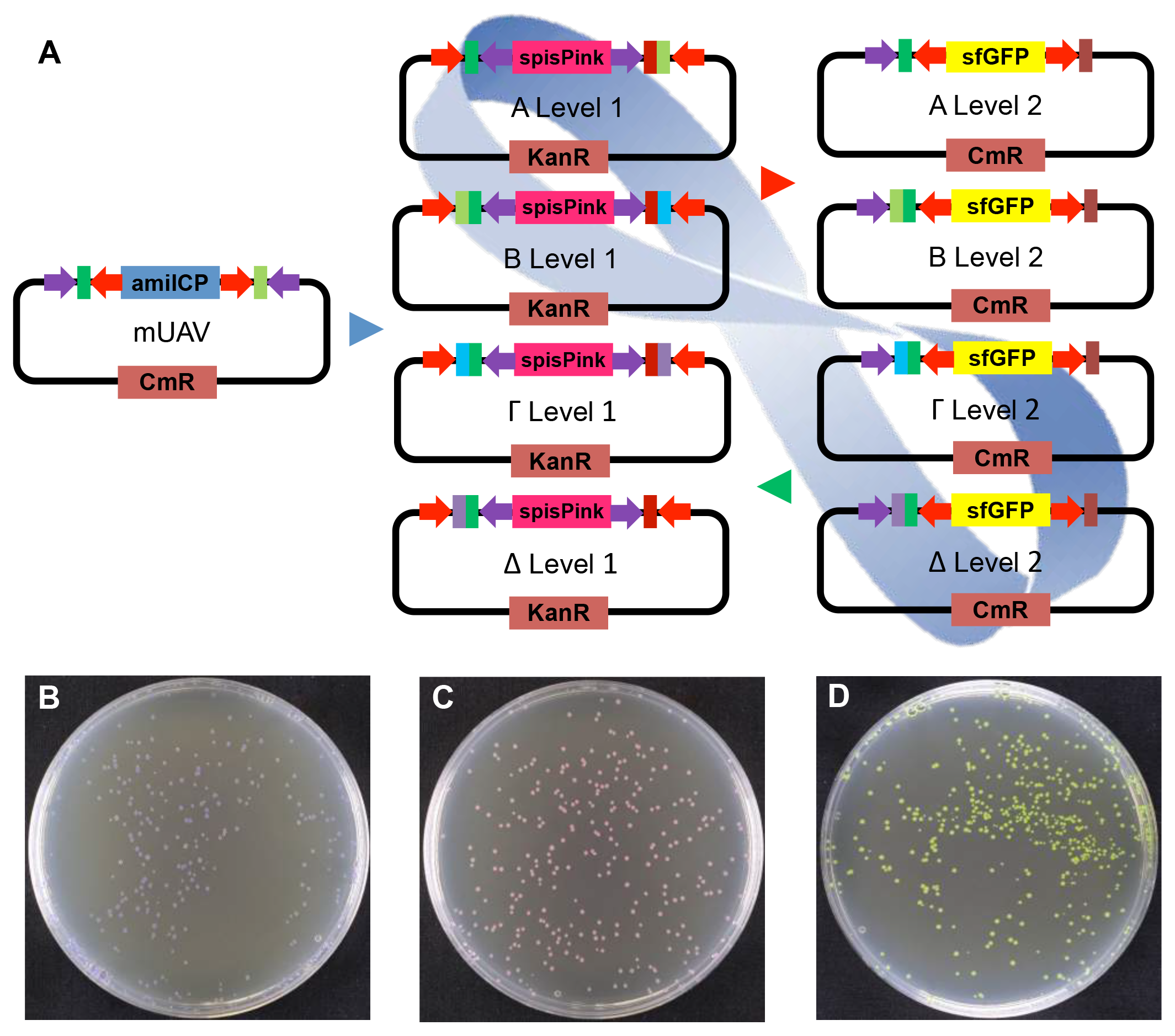
Mobius Assembly framework. A. Mobius Assembly uses a two-level (Level 1 and 2) approach for transcriptional unit (TU) and multi-TU augmentation. Each level is comprised of four Acceptor Vectors. The four Level 1 Acceptor Vectors (A, B, Γ, and Δ) carry *spisPink gene* as the visible cloning selection marker and confer Kanamycin resistance. The four Level 2 Acceptor Vectors (A, B, Γ, and Δ) carry *sfGFP gene* as the visible cloning selection marker and confer Chloramphenicol resistance. The standard parts stored in mUAVs are released and fused in a Level 1 reaction to form a TU. Up to four Level 1 TUs can be fused in a Level 2 reaction to form a multi-TU cassette. Switching back and forth between Level 1 and 2 leads to further expansion of multi-TUs according to the geometric sequence: 1,4, 16, 64, …. Red arrows denote *Aar*I restriction sites and Purple arrows *Bsa*l restriction sites. B, C, and D. *E. coli* colonies carrying mUAV, Level Acceptor 1 Vector A, and Level 2 Acceptor Vector A, which respectively exhibit purple, magenta and yellow colour after overnight incubation. Successful assembly produces white colonies.

**Figure 3.**
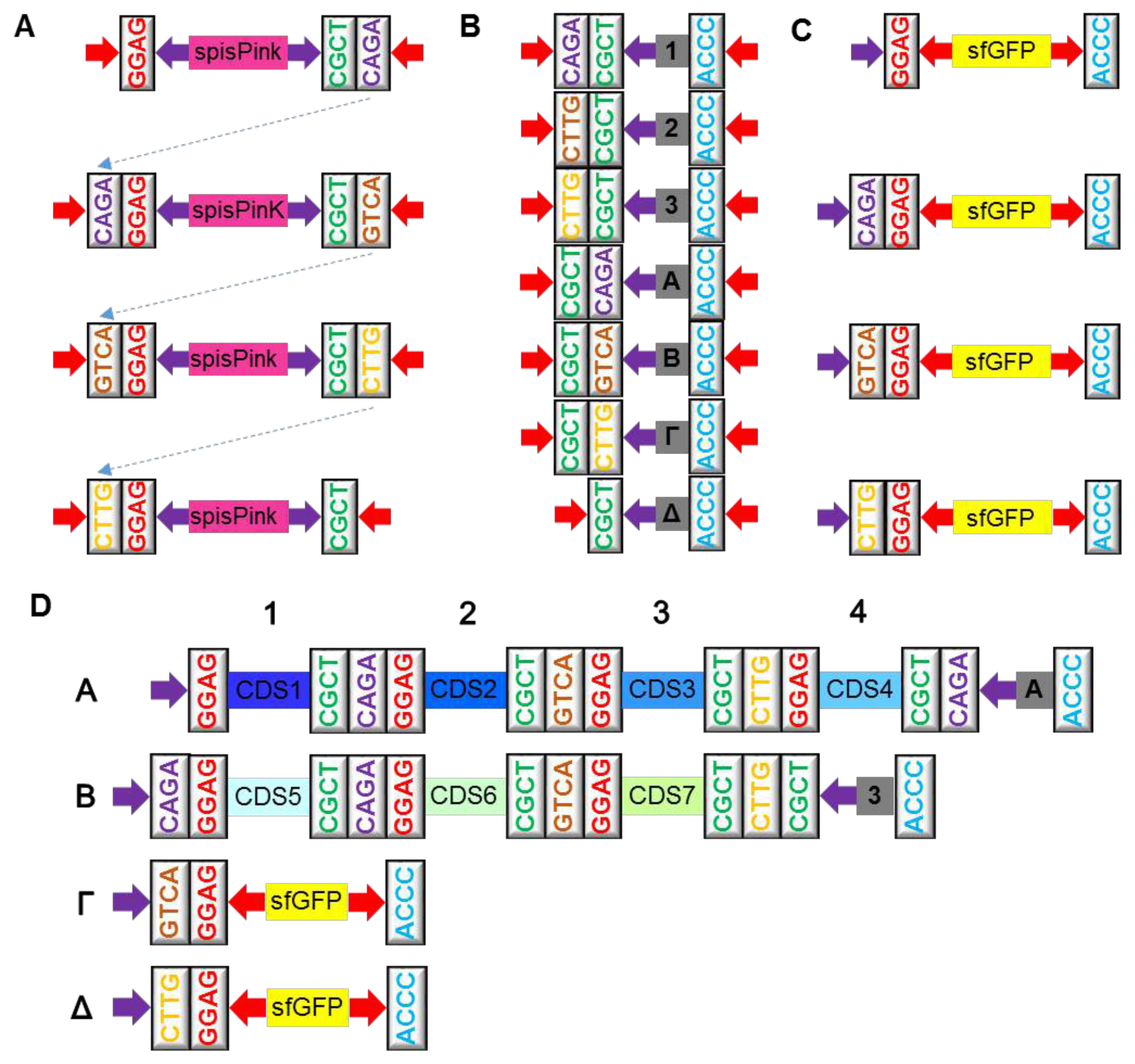
Mobius Assembly vector toolkit. (A) The overhangs of the four Level 1 Acceptor Vectors. *Bsa*l digestion releases the *spisPink* gene upon digestion to expose GGAG and CGCT, between which a TU will be incorporated. Each type of vector has unique overhangs at the 3’ end which guides the assembly of up to four TUs in a Level 2 Acceptor Vector. (B) The overhangs of the four Level 2 Acceptor Vectors. *Aar*l digestion releases the *sfGFP* gene to expose GGAG and ACCC, between which up to four TUs will be fused into with the assistance of an auxiliary plasmid. (C) Seven auxiliary plasmids provide End-to-End linkers and Middle-to-End linkers to assist Level 2 cloning. A, B, Γ and Δ End-to-End linkers provide 5’ and 3’ overhangs and the missing Level 2 overhang when four Level 1 TUs are fused. Middle-to-End linkers 1, 2, and 3 are used when one, two or three Level 1 cassettes are fused in Level 2. They provide 5’ and 3’ overhangs and the CGCT overhang necessary for the cloning back to Level 1. (D) An example of how the auxiliary plasmids are used. A 7-TU construct is generated by combining the four TUs in the Level 2 Acceptor Vector A and the remaining three TUs in Vector B in a Level 1 reaction. Auxiliary Plasmid A is used for the four TUs in Acceptor Vector A, and the Auxiliary Plasmid 3 for the three TUs in Vector B. Red arrows demarcate *Aar*l restriction sites and purple arrows *Bsal* restriction sites.

Mobius Assembly was designed such that the standard parts are released by *Bsa*l digestion, as in MoClo and Golden Braid, to facilitate exchangeability. At the same time, we introduced AarI as a second restriction enzyme to address the domestication issue (Figure 1A). We opted for AarI, because it is a rare cutter that recognizes the 7bp sequence CACCTGC(4/8)^ and leaves a 4bp overhang. Other Type IIS rare cutters leave 2 or 3bp overhangs or contain a large (e.g. 20bp) space between the recognition and cut sites. Golden Braid 2.0^15^ employs three restriction enzymes *Bsa*l, *BsmB*l and *BtgZ*l, all of which recognize 6bp sequences. MoClo^16,17^ also uses 6bp cutters *Bsa*l, *Bpi*l and sometimes *BsmB*l. *Aar*l thus theoretically drops domestication requirements by 75% but still maintains assembly efficiency and a 4bp overhang.

Mobius Assembly embraces the standard 4bp overhangs used by MoClo and Golden Braid. These specific sets of overhangs are becoming more common, being also adopted for Phytobricks, the newly emerging standard part collection for the iGEM registry (Figure 1B). Between MoClo and Golden Braid 2.0, there is partial compatibility of standard parts since they use the same sets of 4bp overhangs and use *Bsa*l for the TU assembly. Full compatibility is possible when a sequence is free from all restricted recognition sites used by each assembly framework. However, because the additional enzymes they require are frequent cutters, direct compatibility is limited. The scarcity of *Aar*l sites facilitates direct use of the available standard parts that have been generated by MoClo or Golden Braid, as well as Phytobricks, by reducing re-domestication requirements. Less domestication also renders Mobius Assembly more efficient for the generation of new standard parts.

To enable theoretically infinite assembly with a simple vector toolkit, we establish a two-level cloning framework that undergo cycled, two-tier hierarchical augmentation, hence the name “Mobius Assembly”. The single TU assembly takes place in Level 1, and multi-TU assembly can be further continued by switching back and forth between Level 1 and Level 2 vectors (Figure 2A). Mobius Assembly enables the assembly or the addition of any number of composite parts as far as the vector or the chassis can handle.

There are four Level 1 Acceptor Vectors, all of which are equipped with a kanamycin resistance gene, since the Mobius Assembly cassette is housed in pSB1 K3 backbone (Figure 2A). In Level 1 vectors, the *spis Pink* chromoprotein gene^25^ serves as a negative cloning selection marker, which colours the colonies pink (Figure 2C); it is released by *Bsa*l digestion and marks colonies with successfully assembled constructs as white. All Level 1 Acceptor Vectors contain the standard fusion sites at the 5’ (GGAG) and 3’ (CGCT) ends to house a (multi-)TU, plus the additional 4bp fusion sites for the cloning of up to four TUs in a Level 2 Acceptor Vector (Figure 3A).

Level 2 is comprised of four Acceptor Vectors, which have the pSB1C3 backbone that confers chloramphenicol resistance (Figure 2A). They all contain the *sfGFP* (strong folded GFP) gene^26^ as a negative cloning selection marker, which makes the colonies yellow (Figure 2D); the selection marker is released by *Aar*l digestion, and successful assembly results in white colonies. Level 2 Acceptor Vectors have the same 5’ overhangs as Level 1 Acceptor Vectors and a common 3’ fusion site (ACCC), where the linkers from the auxiliary plasmids will anneal, providing the appropriate fusion sites to enable the assembly of up to four TUs in a Level 1 Acceptor Vector (Figure 3C).

In total, there are seven auxiliary plasmids that provide four End-to-End and three Middle-to-End 50bp linkers, which confer kanamycin resistance (Figure 3B). In the scenario where four Level 1 vectors are assembled into a Level 2 Acceptor Vector, an End-to-End auxiliary plasmid (A, B, Γ, or Δ) is recruited to provide a linker containing three types of overhangs: i) the 5’ End overhang CGCT, which anneals to the 3’ overhang of Level 1 vector Δ, ii) Level 2 End overhang depending on the type of Level 2 Acceptor Vector (A=CAGA, B=GTCA, Γ=CTTG, or Δ=CGCT) used, and iii) the 3’ End overhang ACCC (Figure 3B and D). The auxiliary plasmid Δ is also used when eight or twelve TUs are assembled into B and Γ Acceptor Vectors, respectively. When less than four TUs are fused together, a Middle-to-End auxiliary plasmid (1, 2, or 3) is used to provide a linker containing three types of overhangs: i) 5’ end overhang depending on the number of TUs being combined (1 = CAGA, 2 = GTCA, or 3 = CTTG), ii) the overhang CGCT necessary to continue assembly back to Level 1, and iii) the 3’ end overhang ACCC (Figure 3B and D). Cloning from Level 2 to Level 1 does not require any auxiliary plasmids. Since the protocol for Golden Gate Assembly with AarI was not optimized before, we tested different ligases and buffers and chose to use T4 ligase and T4 ligase buffer in Level 2 reactions (Supplementary Figure 1). Several thermocycling conditions for restriction digestion and ligation were also tested, but for 4TU Level 2 assembly there were no significant differences (Supplementary Figure 2 and Table 1).

The current most popular Golden Gate variants have either cloning capacity or simplicity. MoClo has high capacity but its vector toolkit is complicated; Level 0 standard parts feed Level 1, which is comprised of seven plasmids^16,17^. Assembly of TUs from Level 1 to multigene constructs can take two directions. The first direction employs seven Level 2 vectors and 21 End Linkers and can assemble up to six TUs per round^16^. The use of seven of the End Linkers results in non-modular constructs, and the cloning cannot further continue to the next level. The exploitation of the other fourteen End Linkers allows the augmentation of up to six TUs. The second direction uses two more levels of vectors, M and P levels, seven for each, each of which contains seven End Linkers^17^. Furthermore, up to six TUs are fused in a Level M vector, while switching between the M and P levels allows multigene augmentation. On the other hand, Golden Braid 2.0 is simple but requires time to reach high capacity^15^. The standard parts feed the α and Ω levels, and the shift between the α and Ω levels doubles the number of TUs.

The design and workflow of Mobius Assembly framework caters to both capacity and simplicity. Having only four Acceptor Vectors in each level and seven auxiliary plasmids, Mobius Assembly vector toolkit is simple. Mobius Assembly framework elevates the assembly capability, in the manner descripted by the exponential geometric sequence a_n_=aır^n-^1 where *a1*=1, *r*=4 and n = 1, 2, 3, 4, and so on. Moreover, addition of new TU (single or multiple) in an already constructed multi-TU is possible by switching between the two cloning levels.

The last feature we have introduced into Mobius Assembly is visible cloning selection by constitutively expressed chromogenic proteins, which replace Blue-White screening with the inducible lacZ operon. To identify effective visible markers, we screened eight chromoproteins (amilCP, amilGFP, spisPink, asPink, aeBlue, mRFP1, and tsPurple) and sfGFP (strong folded GFP), for strong and fast colour development. Expression of these marker genes are controlled by the Anderson promoter J23106 and the rrnBT1-T7TE terminator. *amilCP, spisPink*, and *sfGFP*, which, after overnight incubation, develop strong purple, magenta, and yellow colours, respectively, (Fig 2B-D), were selected as cloning selection markers for Level 0, 1, and 2. The strength and speed of colour development was strain-dependent. After overnight incubation expression for all the chromogenic genes was faster in TOP10 strain than in DH5α; while the colour was clearly identifiable in TOP10, in DH5α it was only possible after longer incubation (e.g. 24 hours). The difference in speed of colour development is probably due to differences in plasmid copy number, as we could extract higher concentrations of the plasmid from TOP10.

The benefits of cloning selection with chromogenic proteins are multifold. By eliminating the need for two expensive chemicals – IPTG and chromogenic substrate (X-gal) – the cloning selection becomes less costly. In addition, cloning chassis are no longer confined to the *E. coli* strains harboring the lacZΔM15 deletion mutation necessary for X-Gal screening. Furthermore, the use of distinct colours in each cloning level assists users in distinguishing between different cloning levels, and can be exploited by automated assembly platforms.

## Proof-of-concept experiments

To validate Mobius DNA Assembly in functionally reconstructing multigene constructs, we assembled genes involved in the violacein and carotenoid biosynthesis as well as chromoprotein TUs. For the proof-of-concept experiment, pigmentation genes were chosen as their colour development facilitated the identification of the correctly assembled constructs with the naked eye.

As the first proof-of-concept experiment for Mobius Assembly, to reconstruct biosynthetic pathways organized into clusters sharing regulatory sequences, four genes from the violacein operon were re-assembled. Violacein is a bisindole pigment mainly produced in bacteria of the genus *Chromobacterium*^27^. The amino acid L-tryptophan, which is colourless within the visible range, is converted to the purple pigment violacein by the sequential activities of five enzymes co-localized in an operon: VioA, VioB, VioC, VioD and VioE (Figure 4A). VioA converts L-tryptophan into indole-3-pyruvic acid imine, which is then dimerized by VioB. VioE catalyzes the conversion of the dimer into protodeoxyviolaceinic acid, which is then converted to protoviolaceinic acid by VioD. The final product, violacein, results from conversion of protoviolaceinic acid via the catalysis by VioC.

**Figure 4.**
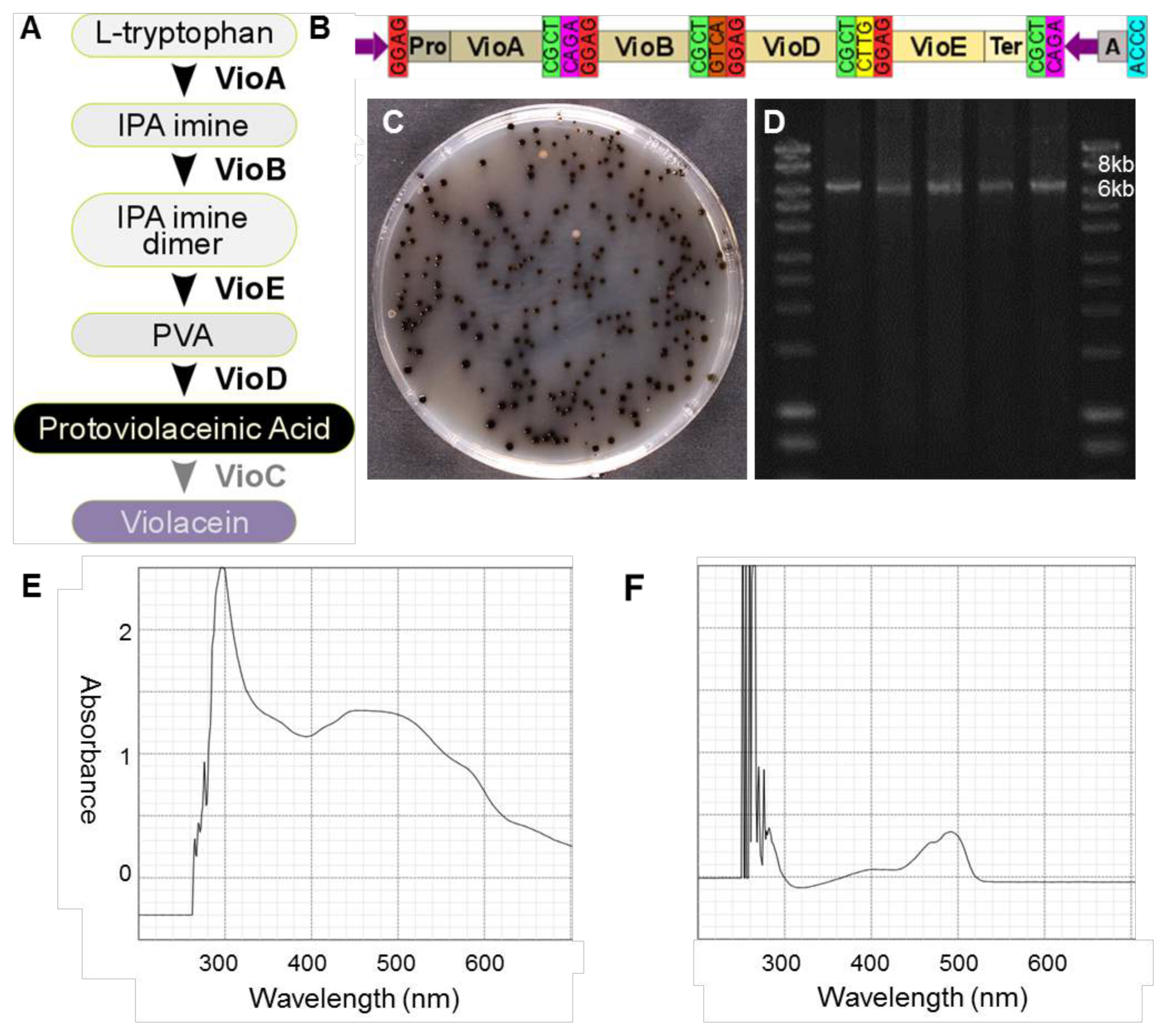
Reconstruction of the violacein biosynthesis operon. (A) Schematic of the violacein biosynthetic pathway showing enzymes mediating the conversion of the intermediates. (B) Diagram showing the assembly of *vioA, vioB, vioD* and *vioE* in a Level 2 Acceptor Vector A. (C) Cells transformed with the *vioABDE* operon form dark green to black colonies due to the production of protoviolaceinic acid. (D) Agarose gel electrophoresis of PCR from five dark purple colonies verified the correct size of the construct (6.5kb). (E) UV-Visible range spectrophotometry of the ethanol extract from a dark green colony showed a wide spectrum of absorbance indicative of protoviolaceinic acid. (F) Spectrophotometry of sfGFP extract from a colony without the recombinant plasmid. IPA: Indole-3-pyruvic acid; PVA: Protodeoxyviolaceinic acid.

We first cloned each of the four coding sequences (*vioA, vioB, vioD* and *vioE*) in the mUAV, to create the standard parts. Next the *vioA* coding sequence was fused to a weak promoter (Anderson Promoter J23103) in Level 1 Acceptor Vector A, *vioB* in Level 1 Acceptor Vector B, *vioD* in Level 1 Acceptor Vector Γ, and *vioE* with the rrnBT1-T7Te terminator in Level 1 Acceptor Vector Δ. All the genes contained their native ribosome binding sites. Finally, the four genes were fused in Level 2 Acceptor Vector A (Figure 4B). The expression of the construct gave colonies with a deep green to black colour due to the production of protoviolaceinic acid, indicating successful reconstruction of the cluster (Figure 4C). Five colonies were selected for colony PCR, which resulted in products of the expected size (Figure 4D). The nature of the pigmentation was identified by spectrophotometry as spanning a wide range of emission wavelengths from UV to the visible spectrum (Figure 4E).

To test the functional reconstruction of a biosynthetic pathway comprised of different TUs, rather than in an operon arrangement, five genes involved in carotenoid biosynthesis were assembled in three different combinations. Carotenoids are group of omnipresent pigments produced by a diverse range of living organisms, including plants, algae, and microbes^28^. *E. coli* cannot naturally synthesize carotenoids; however, introduction of the carotenoid biosynthetic genes results in accumulation of specific variants^29^. The template used in this study is the carotenoid biosynthesis operon from *Pantoea ananatis*^30^. Farnesyl pyrophosphate (FPP), which is a precursor of several isoprenoid compounds, naturally exists in *E. coli* and is the substrate for carotenoid biosynthesis. Figure 5A depicts the carotenoid biosynthesis pathway. In the first step the enzyme geranylgeranyl pyrophosphate (GGPP) synthase encoded by the *crtE* gene takes isopentenyl pyrophosphate (IPP) along with farnesyl pyrophosphate (FPP) to generate GGPP; GGPP is then converted to a carotenoid intermediate, phytoene, via the action of phytoene synthase encoded by *crtB* gene. The first carotenoid, lycopene, exhibits a pink colour, and results from desaturation of phytoene by *crtI*. The lycopene cyclase from the *crtY* gene mediates conversion of lycopene to β-carotene, which is orange in colour. β-carotene is converted to yellow zeaxanthin by the enzyme beta-carotene hydroxylase encoded by *crtZ*.

**Figure 5.**
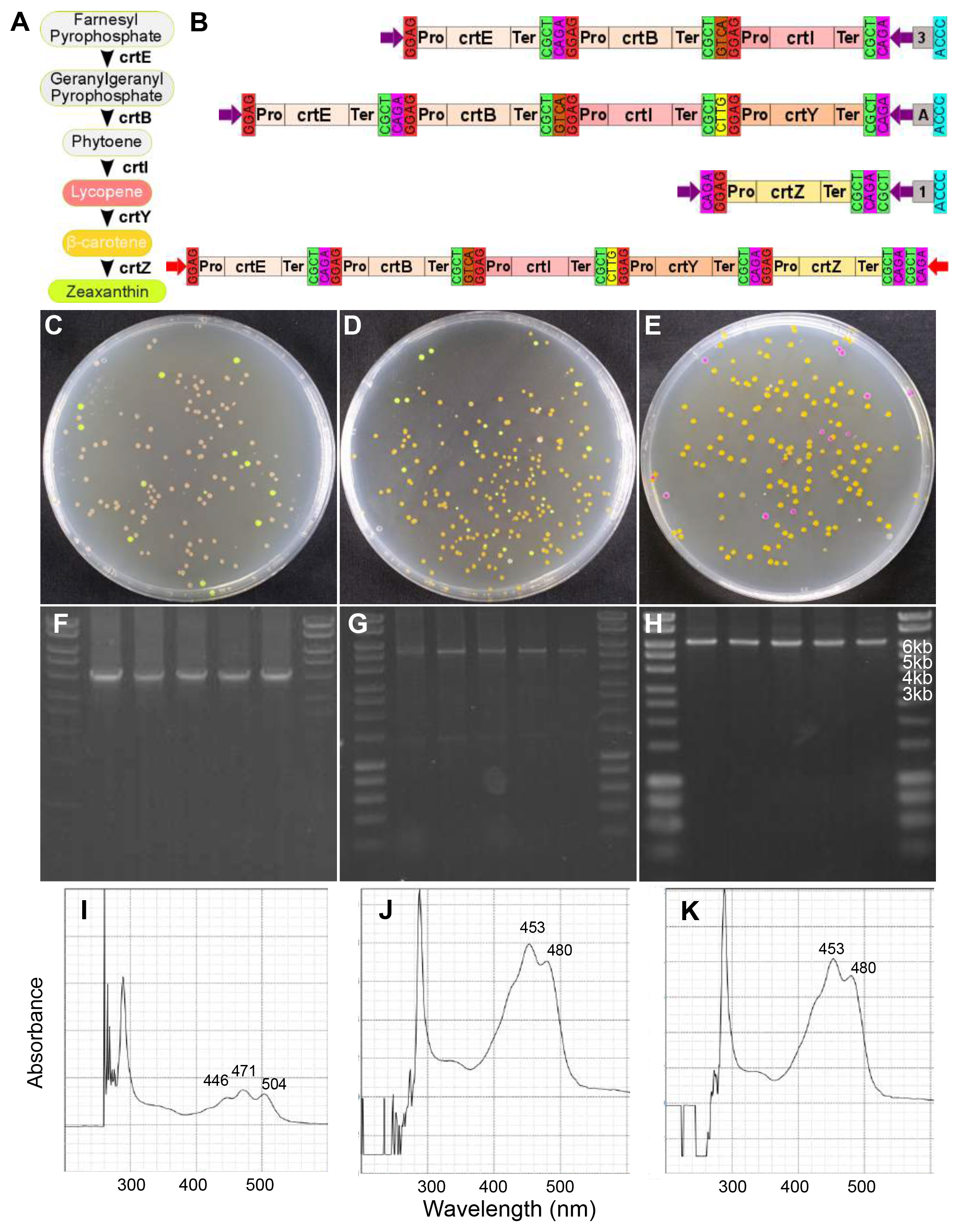
Reconstruction of the carotenoid biosynthesis transcriptional units. (A) A schematic of carotenoid biosynthetic pathway showing the enzymes mediating the production of zeaxanthin as the final product. (B) The multi-TU constructs made by Mobius Assembly to produce lycopene (*crtEBI*) and β-carotene (*crtEBIY)* (in Level 2 Acceptor Vectors) and for zeaxanthin (*crtEBIYZ*) by assembling *crtEBIY* and *crtZ* back in a Level 1 A Vector. Colonies producing lycopene are pink (C), β-carotene orange (D), and zeaxanthin yellow (E). Cells carrying intact Level 2 Vectors produced bright yellow colonies, and Level 1 Vectors pink colonies. Gel electrophoresis of the PCR from five colonies (pink, orange and yellow from each cloning, respectively) verified the correct size of the constructs; 4.3kb for lycopene (F), 5.7kb for β-carotene (G), and 6.7kb zeaxanthin (H). UV-Visible spectrophotometry showed expected peaks for lycopene (446nm, 472nm, and 503nm, I), β-carotene (450nm and 478nm, J) and zeaxanthin (450nm and 478nm, K)^32^.

Cloning of the carotenoid biosynthesis pathways proceeded as follows. Firstly, in Level 0, five genes involved in the carotenoid biosynthesis, *crtE, crtB, crtI, crtY* and *crtZ*, were cloned into the mUAV. Next we assembled them into TUs (promoter:coding_sequence), each of which included the terminator rrnBT1-T7Te. J23110:*crtE* and J23103:*crtZ* were cloned in Level 1 Acceptor Vector A and J23103:*crtB*, J23103:*crtI*, J23103:*crtY* in Level 1 Acceptor Vectors B, Γ, and Δ, respectively. The weak promoter J23103 was chosen after we observed that strong overexpression of these carotenoid biosynthesis genes was lethal to the cells (data not shown). To synthesize lycopene, three TUs *crtE, crtB*, and *crtI* were assembled in a Level 2 Acceptor Vector A (Figure 5B), and successfully constructed cassettes resulted in pink-coloured colonies (Figure 5C). For biosynthesis of β-carotene, four TUs *crtE, crtS, crtI*, and *crtY*were assembled, resulting in orange coloured colonies (Figure 5D). To form the final expression cassette TU *crtZ* was cloned in Level 2 Acceptor Vector B, which was fused with *crtEBIY* in a second assembly step back to Level 1 Acceptor Vector A. This five TU construct led to colonies with a yellow colour, consistent with zeaxanthin accumulation (Figure 5E). The size of the constructs was verified by colony PCR amplification of the insert (Figure 5 F, G, H). The identity of the carotenoid variants produced by each construct was verified using spectrophotometry; the expected emission peaks were observed for lycopene (Figure 5I), β-carotene (Figure 5J) and zeaxanthin (Figure 5K) extracts.

Lastly, we tested the hierarchical assembly capacity of Mobius Assembly by assembling a 16TU construct, which is 18.2kb in size (20.4kb with the vector). To create this high-level multi-TU construct, carotenoid, chromoprotein, and violacein TU modules were combined. The chromoproteins for this experiment were selected such that the individual colours would be detectable, even in combination. Eight chromoprotein genes were cloned into Level 1 Acceptor Vectors, each of which was combined with the weak promoter J23103 and weak transcription terminator for the *E.coli* RNA polymerase, T7Te. We chose to have expression of each gene low to avoid any possible toxicity due to over production of pigments and/or competition for transcription/translation machinary^31^. The chromoprotein TUs were grouped into two categories of four genes each, according to their colour ranges – yellow (*scOrange, amilGFP, amajLime, fwYellow*) and pink (*tsPurple, efforRed, asCP, mRFP1*) – and they were respectively assembled in Level 1 Acceptor Vectors A, B, Γ, and Δ.

The single chromoprotein TUs were then fused in Level 2 Acceptor Vectors to form 4TU constructs. More specifically, the yellow group was assembled in Level 2 Acceptor Vector B, and the pink group in Γ (Figure 6A). The successful constructs were identified firstly by their displayed composite colony colour (yellowish and pink) and secondly by colony PCR of the insert. The violacein operon was reconstructed as described above, but in Level 2 Acceptor Vector Δ. In addition, the carotenoid biosynthesis module *crtEBIY*, as described above, was used in Level 2 Acceptor Vector A. In the final assembly step, the four Level 2 Vectors were fused back to Level 1 Acceptor Vector A to generate the 16TU construct (Figure 6B). Again, the correct assemblies were distinguished by the black colour of the colonies due to dominant pigmentation by protoviolaceinic acid (Figure 6C). Six colonies were selected for DNA plasmid isolation and double restriction digestion with *EcoRl* and *Pstl* or *Pstl* and *Alel*, which resulted in the anticipated patterns of DNA bands (Figure 6D and 6E). The presence of all the 16 TUs in the final construct was further verified by Sanger sequencing.

**Figure 6.**
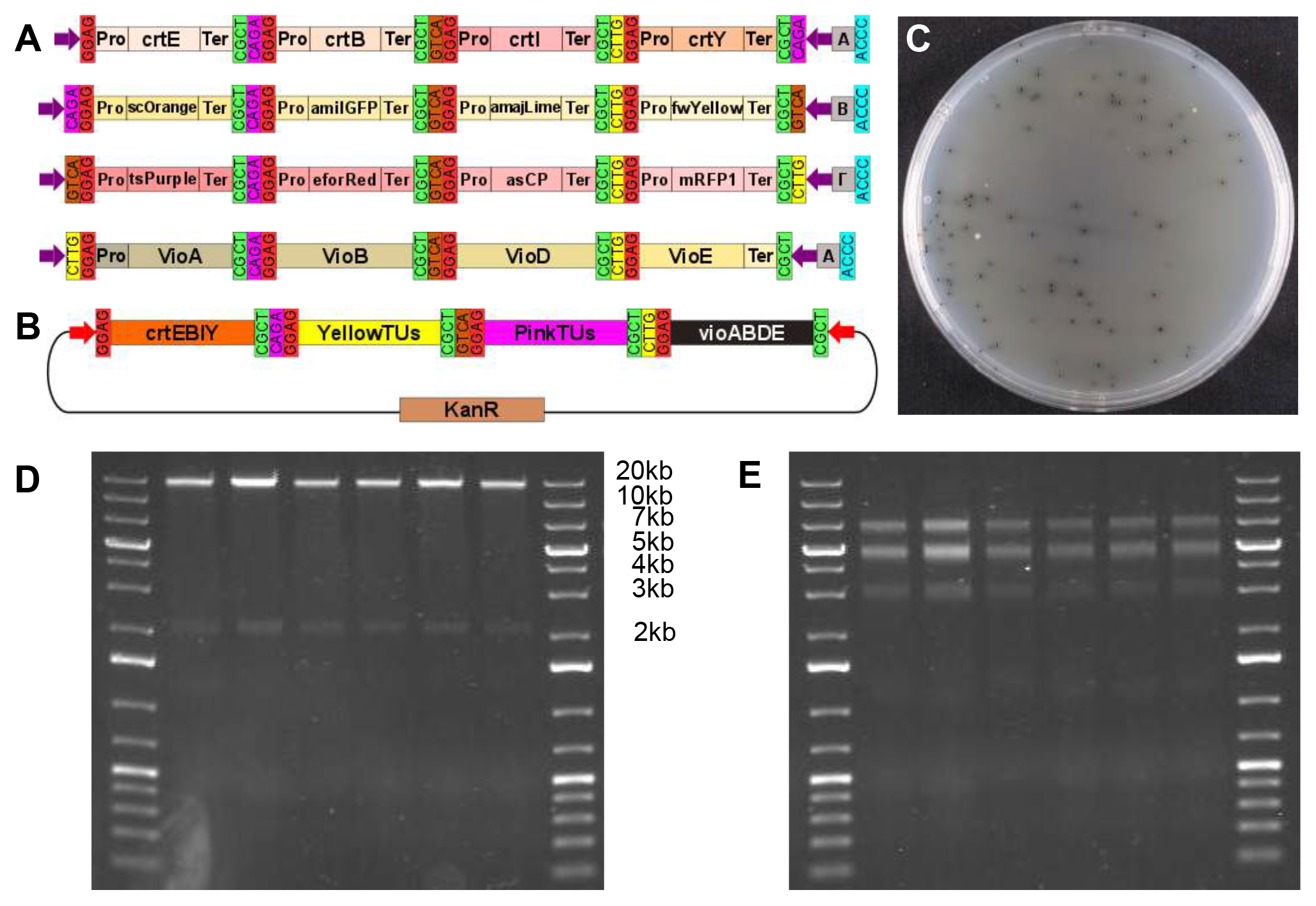
Proof-of-concept assembly of 16TU construct. (A) A schematic showing the four intermediate Level 2 constructs for the assembly of the 16-TU construct. The carotenoid biosynthesis genes *crtE, crtB, crtI*, and *crtY* assembled in the Vector A, the yellow chromoprotein genes *scOrange, amilGFP, amajLime*, and *fwYellow* in the Vector B, the pink chromoprotein genes *tsPurple, eforRed, spisPink*, and *mRFP1* in the Vector Γ, and the violacein biosynthesis genes *vioA, vioB, vioD* and *vioE* in the Vector Δ. (B) A schematic of the 16TU construct derived from the assembly of the four Level 2 cassettes, each containing 4-TUs, in the Level 1 Acceptor Vector A. (C) Cells transformed with the successfully assembled 16TU construct grew into black colonies due to predominant colouring by protoviolaceinic acid. (D) Gel electrophoresis of six plasmids (isolated from the black colonies) digested with *Pst*l and *EcoR*l resulting in bands of expected sizes - 18.3kb for the insert and 2.2kb for the vector. (E) The same plasmids were digested with *Pst*l and *Ale*l resulting in the bands of expected sizes - 7.1kb, 5.3 and 4.9kb (appear merged on the gel), and 3.2kb.

## Materials and Methods

### Cell culturing and plasmid preparation

*E. coli* cells (DH5α or TOP10) were grown in 5ml LB growth medium supplemented with appropriate antibiotics for overnight incubation at 37°C, 230rpm. Plasmids were isolated using Monarch (NEB)/PureYield^TM^ (Promega) Plasmid Miniprep. DH5α (NEB) or TOP10 (Thermo Fisher Scientific) ultracompetent cells were transformed with the constructs as follows: 5μl of the DNA solution was incubated with 50μl of the competent cells on ice for 10min, followed by a heat shock at 42°C for 40s and re-cooled on ice for 10min. SOC medium (400μl) was added, and after 1hr incubation at 37°C, 200rpm, 50μl of the cell suspension was plated on LB agar plates with antibiotic selection. The plates were incubated overnight at 37°C.

### Part and vector generation

pSB1C3 (iGEM DNA distribution) was used as the backbone for mUAV and Level 2 Acceptor Vectors, while pSB1K3 (iGEM DNA distribution) was used for the construction of Level 1 Acceptor Vectors. Bacterial promoters and terminators, as well as the genes for the carotenoid and violacein biosynthetic pathways, were cloned from the iGEM DNA distribution kit. Chromoproteins were kind gifts from the Uppsala iGEM Association. All the standard parts were domesticated for *Aar*l and *Bsa*l when necessary and were cloned into mUAV.

To generate the Mobius Assembly Golden Gate TU cassettes, J23106 promoter, a chromoprotein gene (amilCP for mUAV, spisPink for Level 1 Acceptor Vectors, and sfGFP for Level 2 Acceptor Vectors), and the rrnBT1-T7Te terminator were combined. They were then amplified using Q5 DNA polymerase (NEB) with primers bearing *EcoRl*, *Aarl*, *Bsal*, and *Pstl* restriction sites and 4bp overhangs. Subsequently the pSB1C3/pSB1K3 backbones and the Golden Gate cassettes were digested with *EcoR*l HF and Pstl-HF (NEB) for 20min at 37°C followed by purification with PCR Clean-up kit (Macherey Nagel). Ligation was mediated by T7 ligase (NEB) to construct mUAV, Level 1 and Level 2 Acceptor Vectors. To construct the 50bp linkers in the auxiliary plasmids, a short sequence was PCR amplified from *scOrange* gene using primers that contained the appropriate overhangs and the *Aar*l and *Bsa*l recognition sites. The PCR products were purified and then digested for 2hrs at 37^o^C with *AarI* (Thermo Fisher Scientific). A Level 1 Acceptor Vector A was also digested with *AarI* and ligated to the auxiliary plasmid cassettes with 1μl T7 ligase (NEB) for 20min at RT. The constructs were verified by Sanger sequencing (GATC-Biotech or Edinburgh Genomics).

### Golden Gate Assembly

The DNA assembly was carried out in 10μl reaction comprised of ~50μg Acceptor Vector and twice as many molar of the insert parts, in addition to 1μl 1 mg/ml BSA (NEB), 1 μl T4 ligase buffer (NEB or Thermo Fisher Scientific), 0.5μl *Aar*l (Thermo Fisher Scientific) for cloning in mUAV and Level 2 Acceptor Vectors or *Eco*31l (*Bsa*l) (Thermo Fisher Scientific) for cloning in Level 1 Acceptor Vectors, and 0.5μl T4 ligase (NEB or Thermo Fisher Scientific). For reactions with *Aar*l, extra 0.2μl 50x oligos (0.025 mM) of the enzyme recognition sites were added. The one-tube reaction was incubated in a thermocycler for five times cycles of (37°C for 5min, 16°C for 10min) followed by 5min digestion at 37°C and 5min deactivation at 80°C. For the assembly of the 16TU construct, the reaction was set in 20μl with double amount of buffers and enzymes and the thermocycling conditions were altered to: 40 cycles of (37°C for 2.5min, 16°C for 5 min) followed by 5min digestion at 37°C and 5min deactivation at 80°C.

### Pigment spectrophotometry

Lycopene, β-carotene, zeaxanthin, and protoviolaceinic acid were extracted from 100 ml overnight cultures in LB. Cells were harvested by centrifugation at 2500g for 10min and then resuspended in 2ml 96% ethanol and 2min vortexing for cell lysis. The lysate was then centrifuged for 5min at 14000g, and the supernatant was filtered through a 0.22μm filter (Milipore) and used for spectrophotometry for the UV-visible range wavelengths (200–780nm) using Biowave II (Montreal-biotech).

## Acknowledgements

We thank the Uppsala iGEM Association for kind sharing of their chromoprotein collection. This work was supported by the University of Edinburgh Principal’s Career Development PhD Scholarship (to A. I. A.) and the University of Edinburgh Chancellor’s Fellowship and the Royal Society University Research Fellowship (to N. N.). We thank the members of the Biological Form + Function Lab for their help with improving the manuscript.

